# Dystroglycan is a scaffold for extracellular axon guidance decisions

**DOI:** 10.1101/410944

**Authors:** L. Bailey Lindenmaier, Nicolas Parmentier, Caying Guo, Fadel Tissir, Kevin M Wright

**Affiliations:** Vollum Institute, Oregon Health & Science University, Portland, OR 97239; Institiute of Neuroscience, Universite’ Catholique de Louvain, Brussels, Belgium; Howard Hughes Medical Institute, Janelia Farms Research Campus, Ashburn, VA 20147

**Keywords:** Dystroglycan, Celsr3/Adgrc3, axon guidance, extracellular matrix, internal capsule, commissural axon

## Abstract

Axon guidance requires interactions between extracellular signaling molecules and transmembrane receptors, but how appropriate context-dependent decisions are coordinated outside the cell remains unclear. Here we show that the transmembrane glycoprotein Dystroglycan interacts with a changing set of environmental cues that regulate the trajectories of extending axons throughout the brain and spinal cord. Dystroglycan operates primarily as an extracellular scaffold during axon guidance, as it functions non-cell autonomously and does not require signaling through its intracellular domain. We identify the transmembrane receptor Celsr3/Adgrc3 as a binding partner for Dystroglycan, and show that this interaction is critical for specific axon guidance events *in vivo*. These findings establish Dystroglycan as a multifunctional scaffold that coordinates extracellular matrix proteins, secreted cues, and transmembrane receptors to regulate axon guidance.

## Introduction

During neural circuit development, extending axons encounter distinct combinations of cues and growth substrates that guide their trajectory. These cues can be attractive or repulsive, secreted and/or anchored to cell membranes, and signal through cell surface receptors on the growth cones of axons (Kolodkin and Tessier-Lavigne, 2011). Receptors also recognize permissive and non-permissive growth substrates formed by the extracellular matrix, surrounding cells, and other axons (Raper and Mason, 2010). While many cues and receptors that direct axon guidance have been identified, our understanding of how cues in the extracellular space are organized and interpreted by growing axons is far from complete.

In a previous forward genetic screen for novel mediators of axon guidance, we identified two genes, *Isoprenoid Synthase Domain Containing* (*ISPD*) and *Beta-1,4-glucuronyltransferase 1* (*B4gat1*, formerly known as *B3gnt1*), that are required for the functional glycosylation of the transmembrane protein Dystroglycan (Wright et al., 2012). Dystroglycan is comprised of a heavily glycosylated extracellular α-subunit that is non-covalently linked to its transmembrane β-subunit (Barresi and Campbell, 2006). The mature “matriglycan” epitope on α-Dystroglycan is required for its ability to bind extracellular proteins that contain Laminin G (LG) domains, including Laminins, Perlecan, Agrin, Pikachurin, Neurexin, and Slit (Campanelli et al., 1994; Gee et al., 1994; Ibraghimov-Beskrovnaya et al., 1992; Peng et al., 1998; Sato et al., 2008; Sugita et al., 2001b; Wright et al., 2012; Yoshida-Moriguchi and Campbell, 2015; Yoshida-Moriguchi et al., 2010). The intracellular domain of β-Dystroglycan interacts with the actin binding proteins Dystrophin and Utrophin, and can also function as a scaffold for ERK/MAPK and Cdc42 pathway activation (Batchelor et al., 2007; Ervasti and Campbell, 1993; James et al., 1996; Spence et al., 2004). Therefore, Dystroglycan serves as a direct link between the extracellular matrix (ECM) and pathways involved in cytoskeletal remodeling and filopodial formation, suggesting that it can function as an adhesion receptor to regulate cell motility and migration (Moore and Winder, 2010). However, this has not been examined *in vivo*.

Mutations that result in hypoglycosylation of α-Dystroglycan result in a loss of ligand binding capacity and cause a form of congenital muscular dystrophy (CMD) referred to as dystroglycanopathy. Severe forms of this disorder are accompanied by neurodevelopmental abnormalities including type II lissencephaly, hydrocephalus, brainstem and hindbrain hypoplasia, ocular dysplasia, and white matter defects (Godfrey et al., 2011). We previously found that glycosylated Dystroglycan regulates axon guidance in the developing spinal cord and in the optic chiasm by maintaining the basement membrane as a permissive growth substrate and organizing the extracellular localization of Slit proteins in the floor plate (Clements and Wright, 2018; Wright et al., 2012). However, there are a number of outstanding questions about the role of Dystroglycan in axon guidance: Does Dystroglycan function in axons as an adhesion receptor? Is Dystroglycan required for the formation of other axon tracts in the mammalian nervous system? Does Dystroglycan bind additional LG-domain containing proteins important for axon guidance?

Here, we provide genetic evidence that Dystroglycan operates non-cell autonomously and relies on its extracellular scaffolding function to regulate the development of multiple axon tracts in the spinal cord and brain. We identify a novel interaction between Dystroglycan and Celsr3 (Adgrc3), an LG-domain containing transmembrane receptor that regulates axon guidance in the brain, spinal cord, and peripheral nervous system (Chai et al., 2014; Onishi et al., 2013; Tissir et al., 2005; Zhou et al., 2008). Using genome editing to generate a *Celsr3* mutant that is unable to bind Dystroglycan (*Celsr3*^*R1548Q*^), we show that this interaction is specifically required to direct the anterior turning of post-crossing spinal commissural axons *in vivo*. These results define a novel interaction between Dystroglycan and Celsr3 and establish Dystroglycan as a multifunctional regulator of axon guidance throughout the nervous system via its coordination of multiple ECM proteins, secreted cues, and transmembrane receptors.

## Results

### Dystroglycan functions non-cell autonomously as an extracellular scaffold to guide commissural axons

We have previously shown that loss of glycosylation on Dystroglycan or conditional deletion of *Dystroglycan* throughout the epiblast results in defective axon tract formation in the developing spinal cord and visual system. We found that Dystroglycan is required to maintain the basement membrane as a permissive growth substrate and for the proper localization for the secreted axon guidance cue Slit (Clements and Wright, 2018; Wright et al., 2012). However, we have not tested whether Dystroglycan has a cell-autonomous role in regulating the guidance of spinal commissural axons. Examination of E12 spinal cord sections shows that in addition to its enrichment in the basement membrane, Dystroglycan protein was detected in post-crossing commissural axons (arrows, Figure 1A). In cultured e12 commissural axons, Dystroglycan was seen throughout the axon, including the growth cone (arrows, Figure 1B).

**Figure 1:**
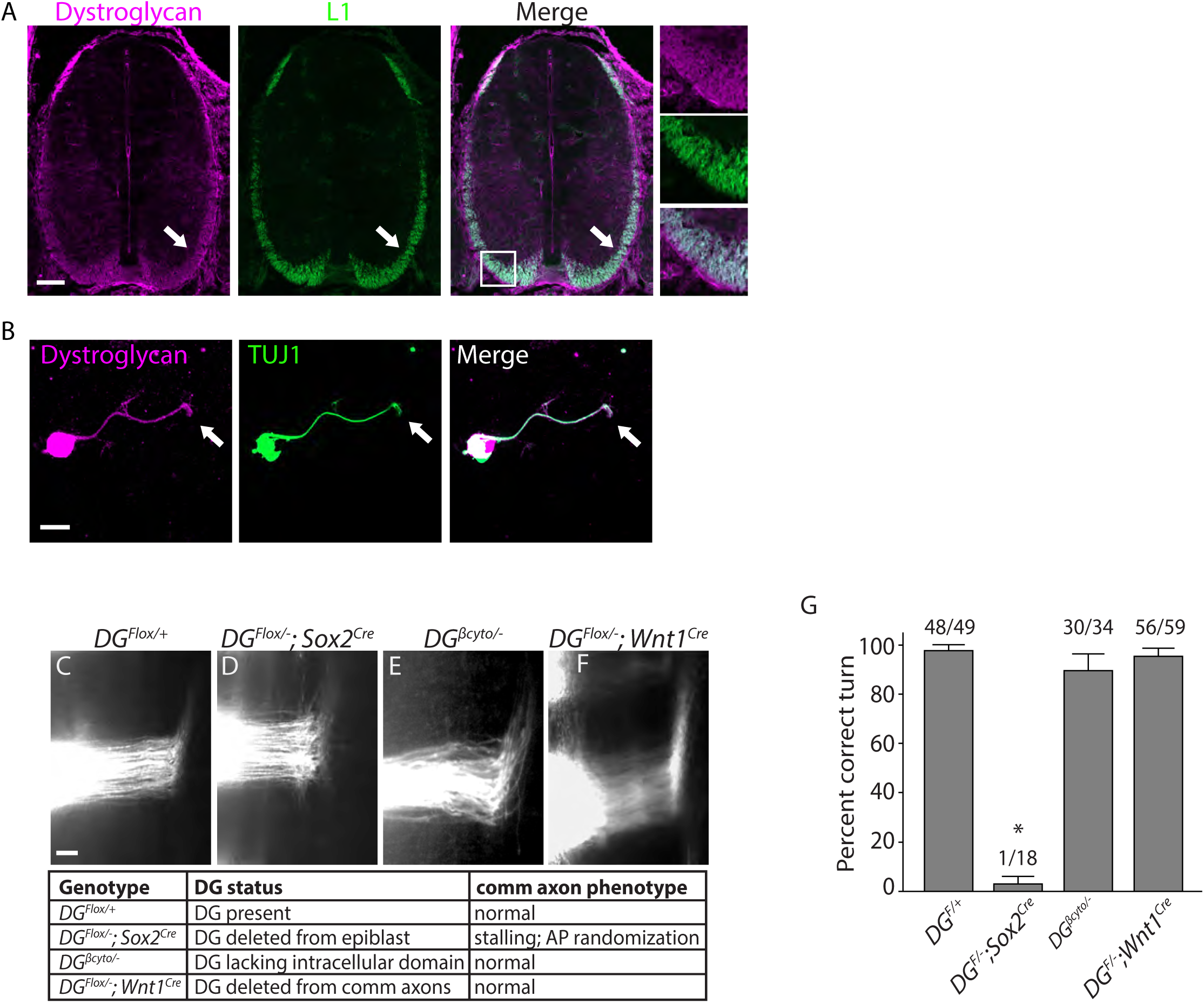
Dystroglycan functions non-cell autonomously to guide spinal commissural axons. **(A)** Immunostaining of E12.5 spinal cords shows Dystroglycan protein (magenta, left panel) expression in post-crossing commissural axons (arrows, L1, green, middle panel). **(B)** Commissural neurons from E12 dorsal spinal cord cultured for two days in vitro (2DIV) were stained with antibodies to Dystroglycan (magenta, left panel), TUJ1 (green, middle panel). Dystroglycan is present throughout the cell body, axon and growth cone (arrow)**. (C-F)** DiI injections in open-book preparations of E12 spinal cords were used to examine the post-crossing trajectory of commissural axons. In controls **(C)**, axons extend through the floor plate, then execute an anterior turn (n=6). In *DG*^*F/-*^*;Sox2*^*Cre*^ mice **(D)**, axons stall within the floor plate and post-crossing axons exhibit anterior-posterior randomization (n=3). **(E)** Commissural axons in mice lacking the intracellular domain of Dystroglycan (*DG*^*βcyto/-*^) show normal crossing and anterior turning (n=3). Conditional deletion of *Dystroglycan* from commissural neurons in *DG*^*F/-*^*;Wnt1*^*Cre*^ mice **(F)** did not affect floor plate crossing or anterior turning (n=8). **(G)** Quantification of open book preparations, with the number of injection sites showing anterior turning over the total number of injection sites indicated above each bar. *p< 0.001, one-way ANOVA, Tukey’s *post hoc* test. Scale bar = 100μm (A), 10μm (B) and 50μm (F-H).

Based on its association with the actin-binding proteins Dystrophin and Utrophin and its ability to regulate filopodial formation *via* ERK/MAPK and Cdc42 activation, we hypothesized that Dystroglycan could function within commissural axons as an adhesion receptor *in vivo*. To test this, we performed DiI labeling in open-book e12.5 spinal cord preparations. In control open book preparations, commissural axons in 48/49 injection sites showed normal floor plate crossing and anterior turning (Figure 1C, G). In agreement with our previous findings, commissural axons in 17/18 of injection sites from mice lacking Dystroglycan throughout the developing spinal cord (*DG*^*F/-*^*;Sox2*^*Cre*^) exhibited stalling within the floorplate and an anterior-posterior (AP) randomization of post-crossing axonal trajectory (*p*>0.001; Figure 1D, G). We next examined commissural axons in which the intracellular domain of Dystroglycan is deleted (*DG*^*βcyto/-*^), rendering it unable to bind dystrophin/utrophin or initiate ERK/MAPK or Cdc42 signaling (Satz et al., 2009). To our surprise, commissural axons in 30/34 injection sites in *DG*^*βcyto/-*^ mutants showed normal floorplate crossing and anterior turning (Figure 1E, G), suggesting that the intracellular domain of Dystroglycan is dispensable for commissural axon guidance. To further test for a cell-autonomous role for Dystroglycan during commissural axon guidance, we examined mice in which *Dystroglycan* is conditionally deleted from commissural axons (*DG*^*F/-*^*;Wnt1*^*Cre*^). 56/59 injection sites in *DG*^*F/-*^*;Wnt1*^*Cre*^ open book preparations displayed normal commissural axon growth and post-crossing anterior turning (Figure 1F, G). Taken together, these results support a model that Dystroglycan functions non-cell autonomously as an extracellular scaffold to guide commissural axons *in vivo*.

### Dystroglycan is required for axon tract development in the forebrain

We next sought to determine whether loss of functional Dystroglycan also affected the formation of axon tracts in other regions of the developing nervous system. We used *ISPD*^*L79*/L79\**^ mutants, which lack glycosylated Dystroglycan, and *DG*^*F/-*^*;Sox2*^*Cre*^ mutants, in which *Dystroglycan* is deleted throughout the epiblast. In both mutant strains we observed severe defects in multiple forebrain axon tracts (Figure 2, Supplemental Figure 1). Abnormalities included fasciculated axons in the upper layers of the cortex (arrows), a large, swirling bundle of axons in the ventral telencephalon (asterisk), and a large axonal projection inappropriately exiting through the ventral diencephalon (arrowheads) (Figure 2A-C).

**Figure 2:**
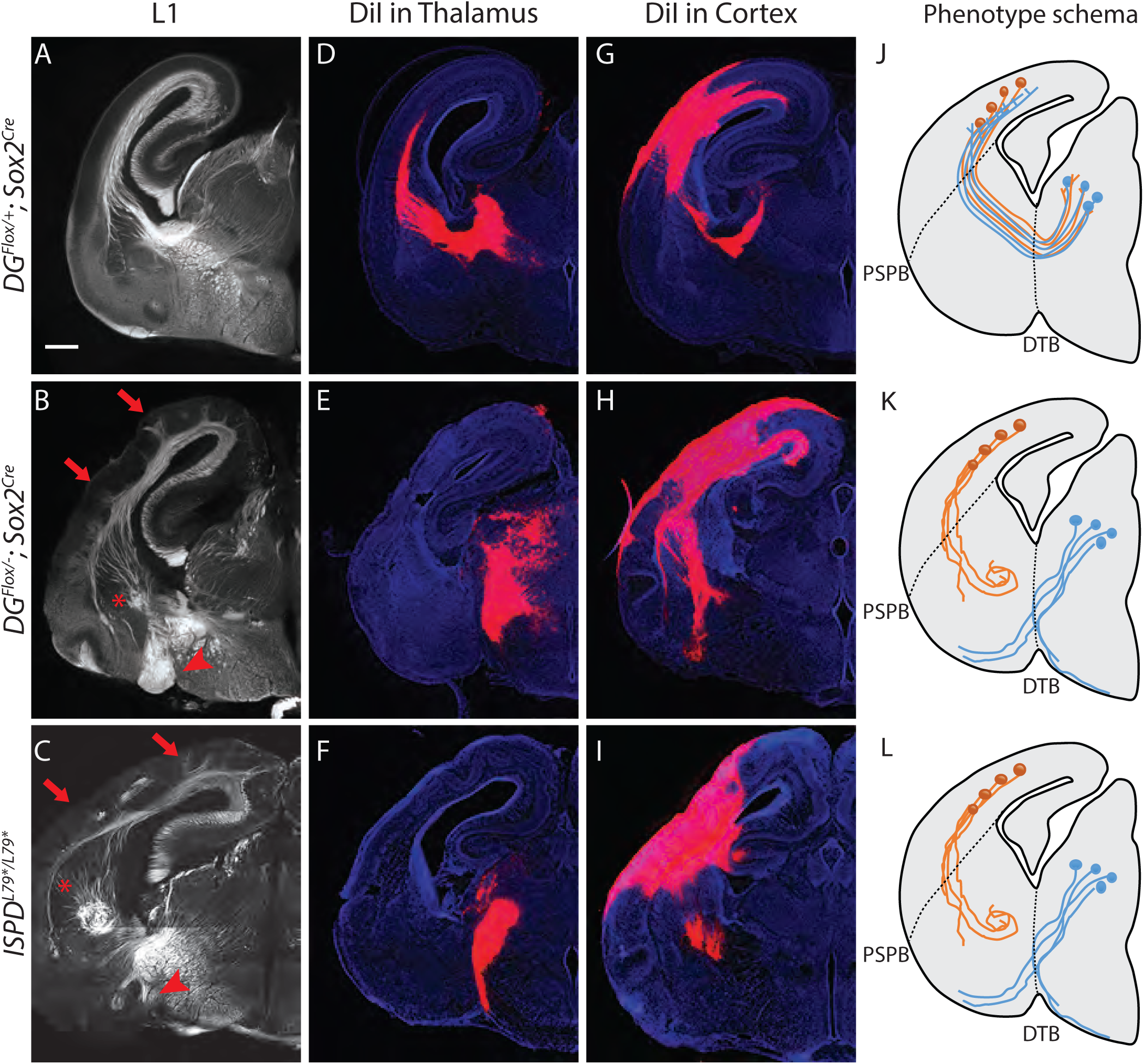
Dystroglycan is required for axon tract formation in the brain. **(A)** L1 immunohistochemistry on P0 brain sections from controls (*DG*^*F/+*^*;Sox2*^*Cre*^) labels descending CTAs and ascending TCAs in the internal capsule. In *DG*^*F/-*^*;Sox2*^*Cre*^ **(B)** and *ISPD*^*L79*/L79\**^ **(C)** mutants, the internal capsule is highly disorganized, with axons projecting into the upper layers of the cortex (red arrows), forming ectopic bundles in the ventral telencephalon (red asterisks), and abnormal projections extending ventrally (red arrowheads). **(D)** DiI injection in the thalamus of controls (*DG*^*F/+*^*;Sox2*^*Cre*^) labels TCAs as they cross the DTB, extend through the ventral telencephalon, across the PSPB, and into the intermediate zone of the cortex. In *DG*^*F/-*^*;Sox2*^*Cre*^ **(E)** and *ISPD*^*L79*/L79\**^ **(F)** mutants, CTAs fail to cross the DTB, and instead project ventrally out of the diencephalon. **(G)** DiI injection in the cortex of controls (*DG*^*F/+*^*;Sox2*^*Cre*^) labels CTAs as they extend across the PSPB, through the ventral telencephalon, and across the DTB into the thalamus. CTAs in *DG*^*F/-*^*;Sox2*^*Cre*^ **(H)** and *ISPD*^*L79*/L79\**^ **(I)** mutants fail to cross the PSPB or take abnormal trajectories through the ventral telencephalon. **(J-L)** Schematic summarizing CTA (brown) and TCA (blue) axon trajectories in controls **(J)**, *DG*^*F/-*^*;Sox2*^*Cre*^ **(K)** and *ISPD*^*L79*/L79\**^ **(L)**. Scale bar = 500μm

To better understand the nature of the axonal defects in *ISPD*^*L79*/L79\**^ and *DG*^*F/-*^*;Sox2*^*Cre*^ mutants, we used anterograde tract tracing. DiI labeling of thalamocortical axons (TCAs) in controls showed that axons cross the diencephalon-telencephalon boundary (DTB), extend dorsolaterally through the ventral telencephalon, and cross the pallial-subpallial boundary (PSPB) before turning medially to extend along the intermediate zone of the cortex (Figure 2D,J). In contrast, TCAs in both *ISPD*^*L79*/L79\**^ and *DG*^*F/-*^*;Sox2*^*Cre*^ mutants largely failed to cross the DTB, and instead extended ventrally out of the diencephalon, often joining the optic tract (Figure 2E,F,K,L). Occasionally, TCAs take a more rostral route through the ventral telecephalon in an abnormal trajectory, where they eventually turn and enter the cortex. These aberrant TCAs then extend into the upper layers of the cortex in large fascicles rather than remaining in the intermediate zone (data not shown).

DiI injections in the cortex of controls labeled corticothalamic axons (CTAs) that project across the PSPB, then execute a ventromedial turn to project through the ventral telencephalon before turning dorsomedially across the DTB into the thalamus (Figure 2G,J). DiI labeling in *ISPD*^*L79*/L79\**^ and *DG*^*F/-*^*;Sox2*^*Cre*^ mutants indicated that many CTAs fail to cross the PSPB. Axons that do cross the PSPB stall or take abnormal trajectories through the ventral telencephalon (Figure 2H,I,K,L). Few CTAs in *ISPD*^*L79*/L79\**^ and *DG*^*F/-*^*;Sox2*^*Cre*^ mutants were able to correctly navigate through the internal capsule to arrive at the thalamus.

In addition to the defects in TCAs and CTAs, other axon tracts within the developing forebrain were malformed in *ISPD*^*L79*/L79\**^ and *DG*^*F/-*^*;Sox2*^*Cre*^ mutants. The anterior commissure was frequently diminished in *ISPD*^*L79*/L79\**^ mutants (Supplemental Figure 1C,D). The lateral olfactory tract (LOT), which contains axons projecting from the olfactory bulb to cortical targets, normally forms directly beneath the pial surface of the ventrolateral rostral forebrain (arrowheads, Supplemental Figure 1A,C). In *ISPD*^*L79*/L79\**^ mutants, the LOT was consistently abnormal, often projecting deeper into the ventrolateral forebrain (arrowheads, Supplemental Figure 1B,D). In contrast, the corpus callosum in *ISPD*^*L79*/L79\**^ mutants appears largely normal, despite the prominent number of axons projecting inappropriately into the upper layers of the cortex (Supplemental Figure 1B,D). Taken together, these results show that glycosylated Dystroglycan is required for proper development of multiple axon tracts in the forebrain.

### Dystroglycan functions non-cell autonomously to guide thalamocortical and corticothalamic axons

Where does Dystroglycan function during forebrain axon tract development? As ascending TCAs and descending CTAs form the internal capsule, they interact with several intermediate targets along their trajectory (Figure 3A, A’). TCAs are guided ventrolaterally across the DTB by Isl1+ guidepost cells, then extend through a permissive “corridor” in the ventral telencephalon formed by lateral ganglionic eminence (LGE) derived cells (Feng et al., 2016; Lopez-Bendito et al., 2006; Metin and Godement, 1996). TCAs contact CTAs at the PSPB, then track along them within the intermediate zone, where they pause for several days before invading the cortical layers (Blakemore and Molnar, 1990; Catalano and Shatz, 1998; Chen et al., 2012). Descending CTAs extend in the opposite direction, first crossing the PSPB, then extending medially through the ventral telencephalon to the DTB along TCAs, where they turn dorsally into the thalamus (De Carlos and O’Leary, 1992; Molnar and Cordery, 1999)

**Figure 3:**
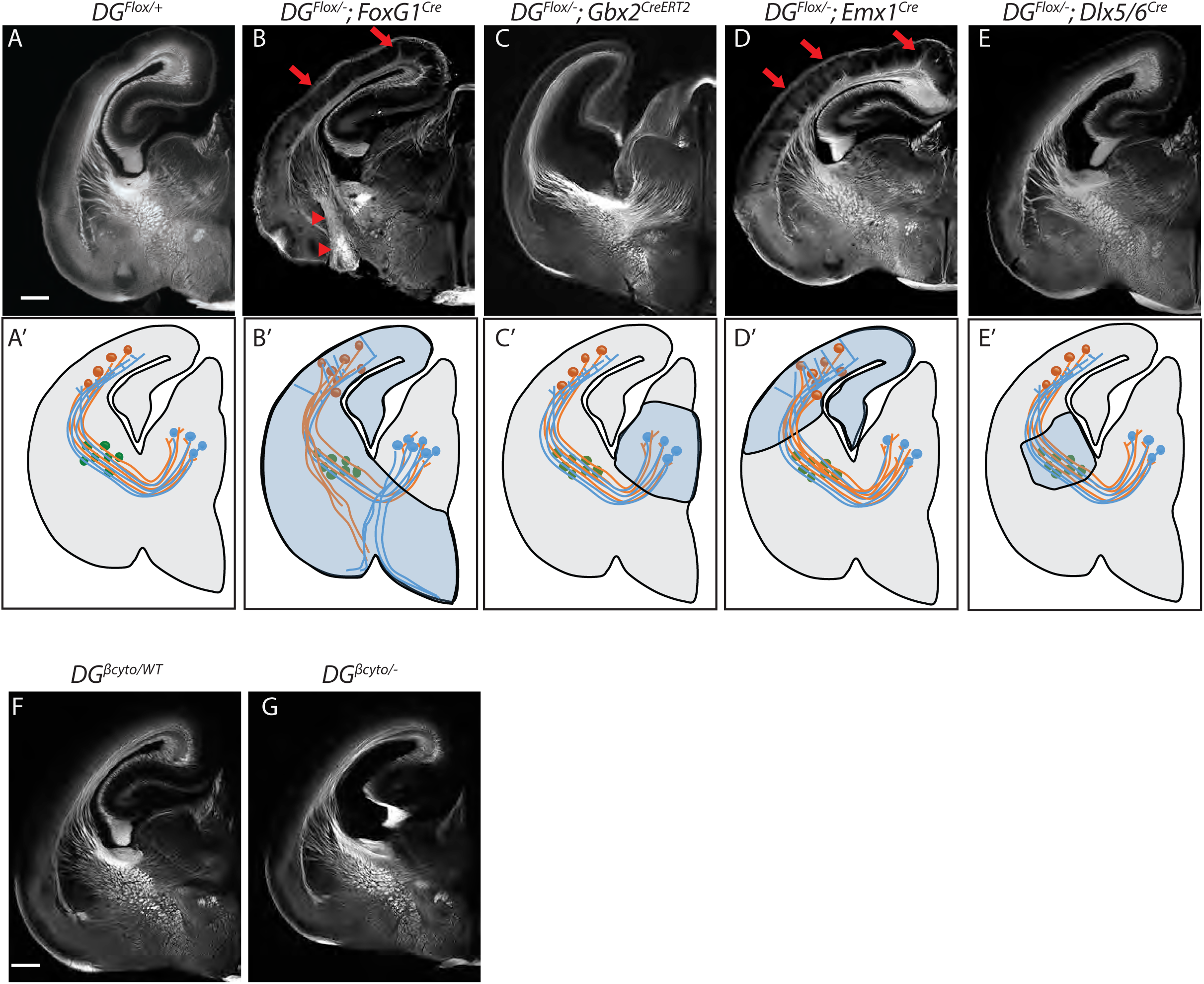
Dystroglycan is required in ventral telencephalon neuroepithelial cells to guide corticothalamic and thalamocortical axons. L1 staining of P0 brain sections from controls (*DG*^*F/+*^) **(A, F)**, *DG*^*F/-*^*;FoxG1*^*Cre*^ **(B)**, *DG*^*F/-*^*;Gbx2*^*CreERT2*^ **(C)**, *DG*^*F/-*^*;Emx1*^*Cre*^ **(D)**, *DG*^*F/-*^*;Dlx5/6*^*Cre*^ **(E),** and *DG*^*βcyto/-*^ **(G). A’-E’** illustrate the recombination patterns in each Cre/CreERT2 line. Deletion of *Dystroglycan* throughout the neuroepithelium of the dorsal and ventral telencephalon in *DG*^*F/-*^*;FoxG1*^*Cr*e^ mutants **(B, B’)** results in abnormal projections in the internal capsule (red arrowheads) and abnormal axonal projections into the upper layers of the cortex (red arrows). Deletion of *Dystroglycan* from the neuroepithelium of the dorsal telencephalon with *Emx1*^*Cre*^ mutants **(D)** results in abnormal axonal projections into the upper layers of the cortex (red arrows), but normal internal capsule formation. Deletion of *Dystroglycan* from the thalamus with *Gbx2*^*CreERT2*^**(C)** or “corridor” cells with *Dlx5/6*^*Cre*^ **(E)** did not affect axon guidance. Deletion of the intracellular domain of Dystroglycan in *DG*^*βcyto/-*^ mutants **(G)** did not affect formation of the internal capsule compared to control littermates **(F)**. A-G Scale bar = 500μm

To identify the specific cellular population in which Dystroglycan is required during internal capsule formation, we took advantage of *Dystroglycan* conditional mutants. We first examined *DG*^*F/-*^*;FoxG1*^*Cre*^ mutants, in which Dystroglycan is deleted in neuroepithelial cells and their progeny throughout the dorsal and ventral telencephalon, but not the developing thalamus (Figure 3B’). Using immunostaining and DiI labeling, we found that both TCAs and CTAs took abnormal trajectories in *DG*^*F/-*^*;FoxG1*^*Cre*^ mutants that were similar to those observed in *DG*^*F/-*^*;Sox2*^*Cre*^ and *ISPD* mutants (arrowheads, Figure 3B, Supplemental Figure 2B, 2B’). To test whether Dystroglycan functions within TCAs, we utilized *DG*^*F/-*^*;Gbx2*^*CreERT2*^ mutants, in which tamoxifen administered at E10 results in recombination throughout the developing thalamus (Supplemental Figure 3A). Based on L1 staining and DiI labeling of TCAs and CTAs, the internal capsule is normal *DG*^*F/-*^*;Gbx2*^*CreERT2*^ mutants (Figure 3C, Supplemental Figure 2C, 2C’). Taken together, these results suggest that Dystroglycan is required in the telencephalon and not in TCAs during internal capsule formation

To further dissect the role of Dystroglycan in the telencephalon, we examined *DG*^*F/-*^*;Emx1*^*Cre*^ mutants in which recombination occurs in neuroepithelial cells and their progeny in the dorsal but not ventral telencephalon (Figure 3D’, Supplemental Figure 3B). *DG*^*F/-*^*;Emx1*^*Cre*^ mutants exhibit significant cortical lamination defects, consistent with the known role of Dystroglycan in regulating cortical migration by maintaining integrity of the neuroepithelial scaffold (data not shown)(Moore et al., 2002; Myshrall et al., 2012; Satz et al., 2008). Despite the abnormal cell body positioning of deep layer neurons, their ability to extend axons across the PSPB, through the internal capsule, and into the thalamus appeared unaffected (Figure 3D, Supplemental Figure 2D). These results suggest that Dystroglycan is not required in CTAs during internal capsule formation. Reciprocal projections from TCAs were likewise able to extend normally through the internal capsule in *DG*^*F/-*^*;Emx1*^*Cre*^ mutants, but upon entering the cortex, formed fasciculated bundles that projected into the upper levels of the cortex (Figure 3D, arrows, Supplemental Figure 2D’). We interpret this as a secondary effect of the migration defects in the cortex *DG*^*F/-*^*;Emx1*^*Cre*^ mutants, as it resembles phenotypes seen when subplate neurons are mislocalized to the upper layers of the cortex (Molnar et al., 1998; Rakic et al., 2006). Finally, we examined the effect of deleting Dystroglycan with *Dlx5/6*^*Cre*^, which recombines in LGE-derived postmitotic neurons in the ventral telecephalon, including those that form the “corridor” (Figure 3E’). CTAs and TCAs in *DG*^*F/-*^*;Dlx5/6*^*Cre*^ mutants extended through the internal capsule and into their target regions normally (Figure 3E, Supplemental Figure 2E, 3E’).

We also tested whether forebrain axon guidance required signaling through the intracellular domain of Dystroglycan. L1 staining shows that the internal capsule, anterior commissure, lateral olfactory tract, and corpus callosum were all normal in *DG*^*βcyto/-*^ mutants (Figure 3G, data not shown), demonstrating that intracellular signaling by Dystroglycan is completely dispensable for forebrain axon guidance. Collectively, we conclude that Dystroglycan is not required in CTAs (*Emx1*^*Cre*^), TCAs (*Gbx2*^*Cre*^), or corridor cells (*Dlx5/6*^*Cre*^), but is required in neuroepithelial cells in the ventral telencephalon (*FoxG1*^*Cre*^). Taken together with our results in spinal commissural axons, these data support a model in which Dystroglycan functions non-cell autonomously as an extracellular scaffold to guide axon tract formation in multiple CNS regions.

### Dystroglycan binds to the axon guidance receptor Celsr3

What are the relevant binding partners for glycosylated Dystroglycan during axon guidance? Dystroglycan binds Laminins to regulate the integrity of basement membranes, which can serve as a permissive growth substrate for extending axons (Clements et al., 2017; Clements and Wright, 2018; Wright et al., 2012). Dystroglycan also binds to the LG domain of Slits to regulate their extracellular distribution in the spinal cord (Wright et al., 2012). *Slit1;Slit2, Slit1;Slit2;Slit3* and *Robo1;Robo2* mutants display defects in commissural axon crossing, as well as internal capsule, anterior commissure, and lateral olfactory tract formation (Bagri et al., 2002; Fouquet et al., 2007; Jaworski et al., 2010; Long et al., 2004; Lopez-Bendito et al., 2007). However, *Slit* and *Robo* mutants do not display the prominent AP randomization seen in the commissural axons of *ISPD*^*L79*/L79\**^ and *DG*^*F/-*^*;Sox2*^*Cre*^ mutants, raising the possibility that Dystroglycan interacts with additional molecules during axon guidance.

We therefore focused our attention on the transmembrane receptor Celsr3/Adgrc3, a mammalian orthologue of the *D. melanogaster* planar cell polarity protein (PCP) Flamingo. Celsr3 is a member of the adhesion GPCR family of proteins and contains two LG domains in its large extracellular region (Figure 4A). *Celsr3*^*-/-*^mutants also show remarkably similar axon guidance defects to *ISPD*^*L79*/L79\**^ and *DG*^*F/-*^*;Sox2*^*Cre*^ mutants, exhibiting AP randomization of post-crossing commissural axons in the spinal cord, as well as defects in anterior commissure and internal capsule formation in the forebrain (Onishi et al., 2013; Tissir et al., 2005; Zhou et al., 2008). We therefore tested whether Dystroglycan could bind to the isolated LG domains of Celsr3. We found that Fc-tagged Dystroglycan bound specifically to Alkaline Phosphatase-tagged Celsr3-LG1 domain (AP-Celsr3-LG1), but surprisingly not the AP-Celsr3-LG2 domain (Figure 4B). Similarly, using tagged Celsr3-LG domains as bait, we found that AP-Celsr3-LG1, but not AP-Celsr3-LG2, was able to bind endogenous glycosylated Dystroglycan from brain lysate (Figure 4C). In a live cell binding assay, AP-Celsr3-LG1 and AP-Slit-C-terminal domains bound to Cos7 cells overexpressing Dystroglycan, whereas AP-Celsr3-LG2 did not (Figure 4D). These results identify Celsr3, via its LG1 domain, as a novel binding partner for Dystroglycan.

**Figure 4:**
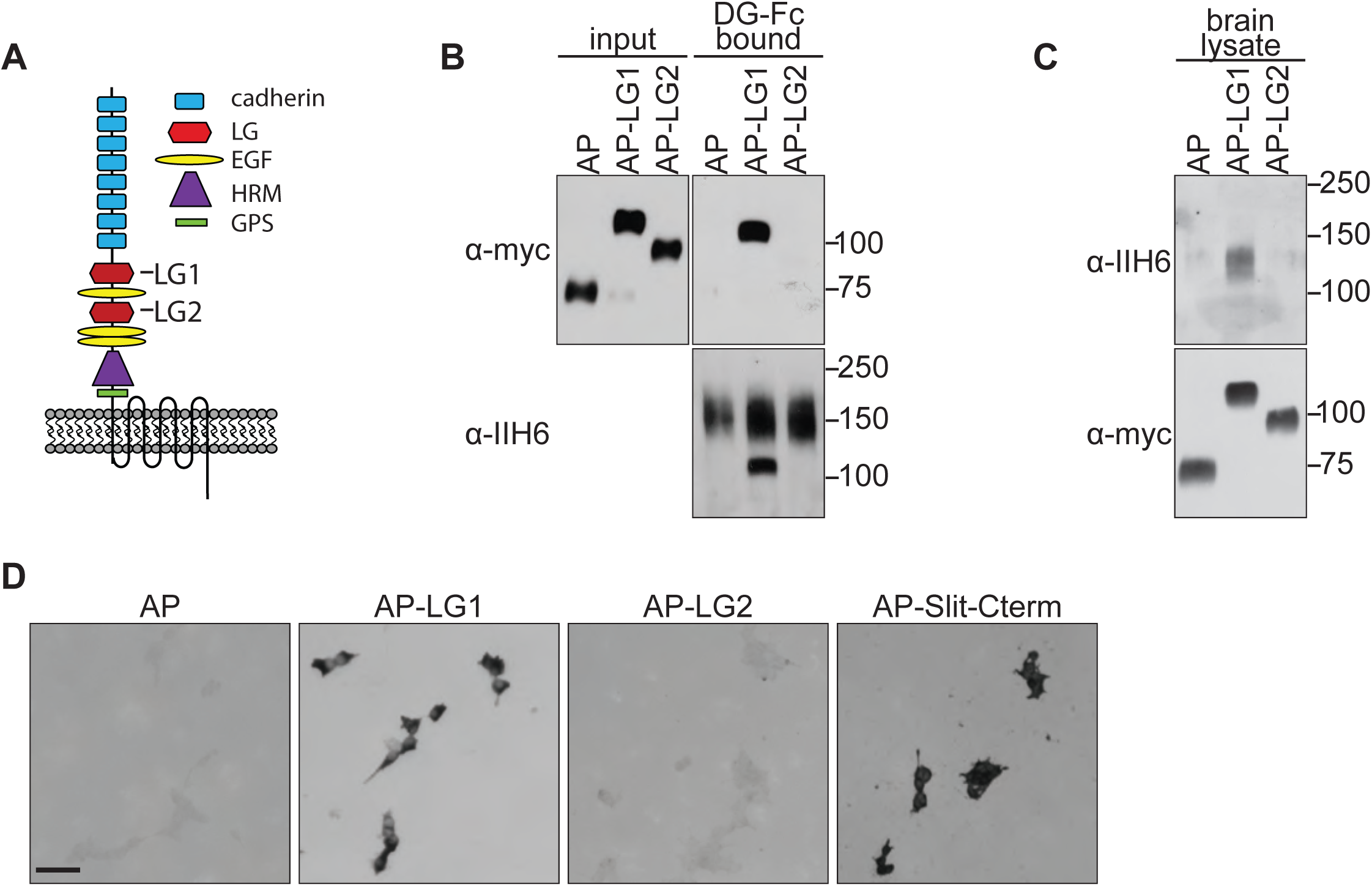
Dystroglycan interacts with the LG1 domain of Celsr3. **(A)** Schematic of Celsr3 protein structure, highlighting the location of Cadherin, Laminin G (LG), EGF, Hormone Receptor Domain (HRM) and GPCR Proteolytic Site (GPS) domains. **(B)** Fc-tagged α-Dystroglycan (Fc-DG) secreted from 293T cells was incubated with Alkaline Phosphatase (AP)-tagged Celsr3-LG1, Celsr3-LG2, or AP-tag alone, and complexes were isolated on Protein A/G beads. DG-Fc interacts selectively with Celsr3-LG1, but not Celsr3-LG2. **(C)** AP-Celsr3-L1, AP-Celsr3-LG2, or AP-tag alone were incubated with WGA enriched brain lysate, and complexes were purified with Ni-NTA beads. AP-Celsr3-LG1 binds endogenous glycosylated Dystroglycan, whereas AP-Celsr3-LG2 and AP-tag do not. **(D)** COS7 cells transfected with full-length Dystroglycan were incubated with AP-tag, AP-Celsr3-LG1, AP-Celsr3-LG2, or AP-Slit-Cterm. Both AP-Celsr3-LG1 and AP-Slit-Cterm exhibited selective binding. Scale bar = 50μm

We next sought to better understand why Dystroglycan binds to Celsr3-LG1, but not Celsr3-LG2. Recent crystal structures have provided insight into how the glycan chains of Dystroglycan bind specifically to LG domains (Briggs et al., 2016). GlcA-Xyl repeats (matriglycan) on Dystroglycan bind a groove in the Laminin-α2-LG4 domain that contains a Ca^2+^ binding site surrounded by several basic residues and a glycine at the tip of the loop (Figure 5A). These residues are all present in the LG domains of the known Dystroglycan binding proteins Laminin-α1, Agrin, Perlecan, Pikachurin, Neurexin, and Slit, suggesting they represent a conserved binding motif between LG domains and the glycan chains on Dystroglycan. Alignment of Celsr3-LG1 with Laminin-α2-LG4 shows significant sequence similarity, including the conservation of the basic residues, the Ca^2+^ binding site, and the glycine at the end of the loop (Figure 5A). This region of Celsr3-LG1 is also evolutionarily conserved (Supplemental Figure 4A). In contrast to Celsr3-LG1, Celsr3-LG2 lacks a Ca^2+^ binding site, the basic residues, and the glycine, and exhibits no sequence conservation with other Dystroglycan-binding LG domains (data not shown), likely explaining its lack of binding.

**Figure 5:**
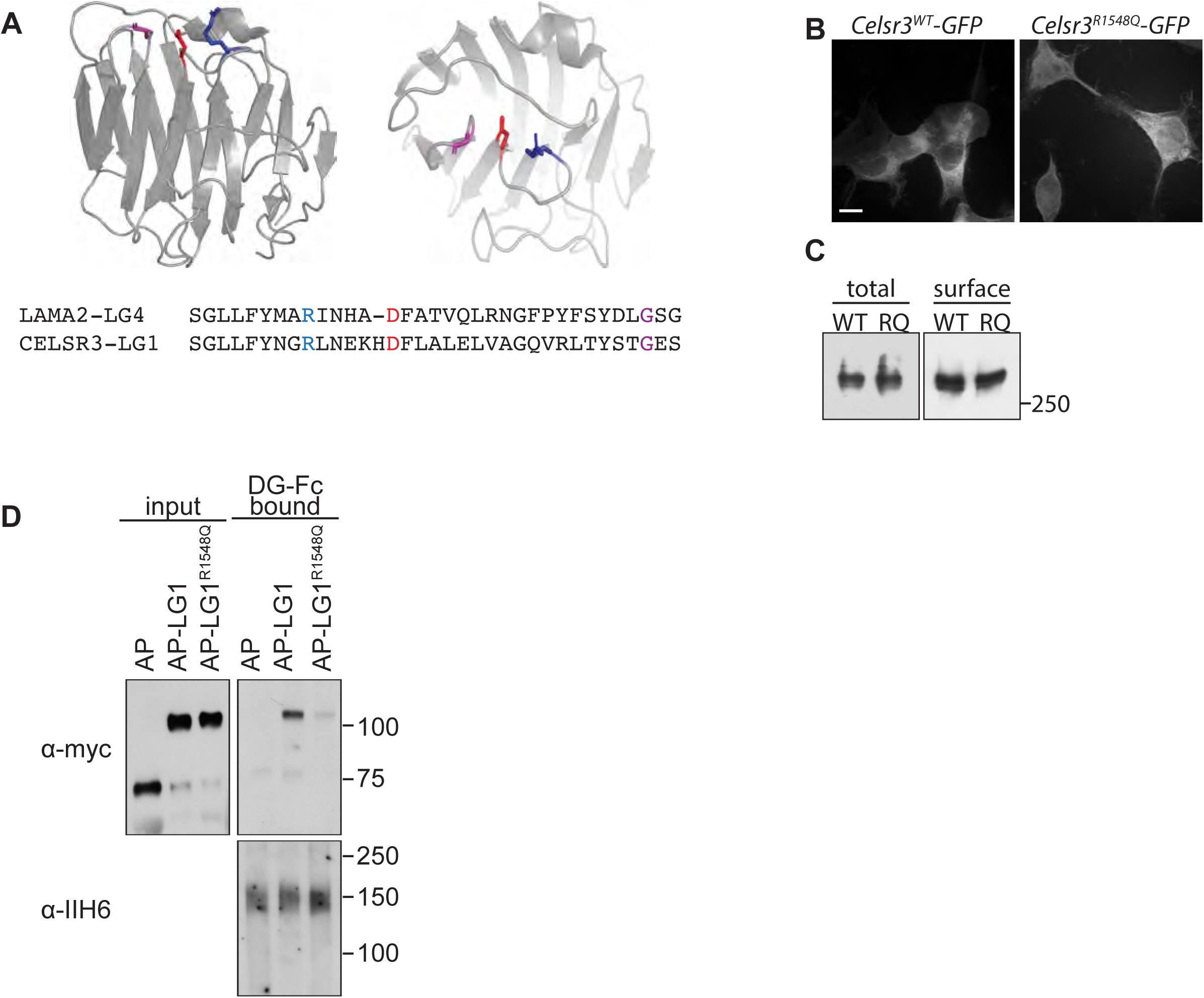
Dystroglycan binding requires specific motifs in Celsr3 LG1. **(A)** Top: schematic showing the structure of the LG4 domain of Laminin-α2 (PDB:1OKQ), highlighting conserved residues critical for Dystroglycan binding: Argenine2803 (blue), Aspartate2808 (red) and Glycine2826 (purple). Bottom: Partial sequence alignment of murine Celsr3-LG1 (amino acids 1540-1574) with murine Laminin-α2-LG4 (amino acids 2795-2828) shows conservation at Argenine1548 (blue), Aspartate1564 (red) and Glycine1572 (purple) of Celsr3. **(B-C)** 293T cells transfected with Celsr3-GFP or mutant Celsr3^R1548Q^-GFP showed no differences in expression levels or cell surface localization by immunocytochemistry **(B)** or western blotting **(C)**. **(D)** Mutation of Celsr3-LG1 at Argenine1548 (AP-LG1^R1548Q^) results in loss of binding to FC-tagged Dystroglycan. Scale bar = 10μm.

To test whether this conserved region of Celsr3-LG1 was required for Dystroglycan binding, we generated GFP-tagged Celsr3 with a mutation at position 1548 (Celsr3^R1548Q^-GFP). This residue corresponds to R2803 in the LG4 domain of Laminin-α2, and is required for its binding to glycosylated Dystroglycan (Wizemann et al., 2003). Compared to wild-type Celsr3-GFP, in which the C-terminal 346 amino acids of the intracellular domain of Celsr3 were replaced with the coding sequence for EGFP, Celsr3^R1548Q^-GFP showed similar subcellular localization and both total and cell-surface expression in 293 cells, suggesting that the R1548Q mutation does not affect the folding or stability of Celsr3 (Figure 5B-C). We next investigated how mutating this residue in the isolated LG1 domain (AP-Celsr3-LG1^R1548Q^) would affect binding to Dystroglycan. Compared to wild-type AP-Celsr3-LG1, AP-Celsr3-LG1^R1548Q^ exhibited markedly reduced binding to DG-Fc, indicating that the conserved binding interface is critical for the specificity of this interaction (Figure 5D).

### Dystroglycan:Celsr3 interactions are specifically required for anterior turning of commissural axons

The axon guidance phenotypes we observed in *Dystroglycan* and *ISPD* mutants are similar to those seen in *Slit/Robo* and *Celsr3* mutants. However, because Dystroglycan binds multiple LG-domain containing proteins through its glycan chains, the phenotypes identified in *Dystroglycan* and *ISPD* mutants likely reflect interactions with multiple extracellular proteins, including Laminins, Slits and Celsr3. To define which aspects of Dystroglycan-dependent axon tract formation require interactions with Celsr3, we used CRISPR/Cas9 genome editing to generate a knock-in mouse carrying an arginine-to-glutamine mutation at position 1548 in Celsr3 (*Celsr3*^*R1548Q*^). *Celsr3*^*R1548Q/R1548Q*^ mice are viable and fertile, as opposed to *Celsr3*^*-/-*^ mice, which die immediately after birth due to respiratory defects (Tissir et al., 2005). Analysis of brain lysates indicated that Celsr3 protein in *Celsr3*^*R1548Q/R1548Q*^ mice migrates at the correct molecular weight, is present at normal levels, and does not lead to compensatory changes in the levels of Celsr1 (Figure 6A).

**Figure 6:**
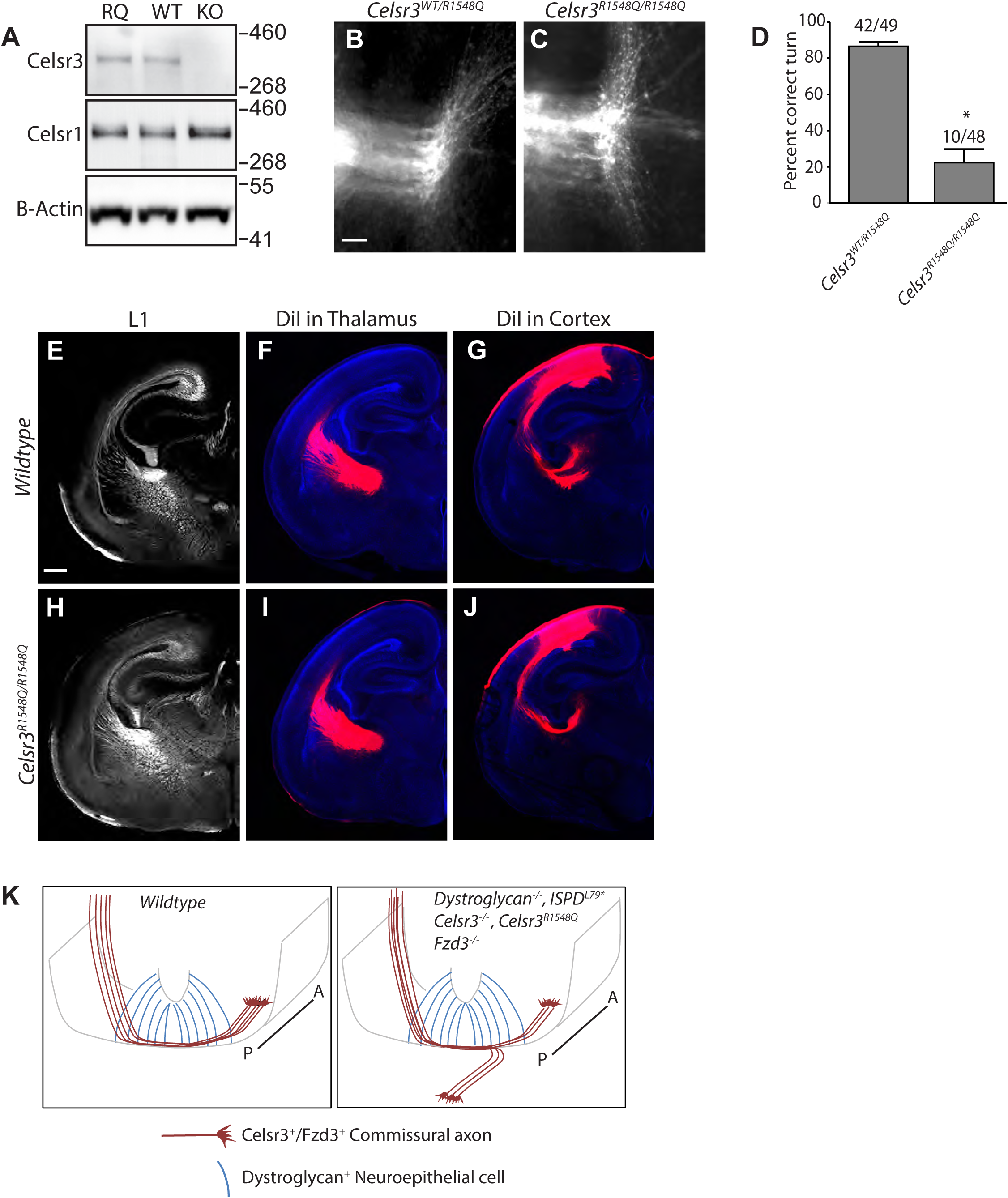
Dystroglycan:Celsr3 interactions are required for spinal commissural axon guidance. **(A)** Western blotting of brain lysates from *Celsr*^*R1548Q/R1548Q*^ mutants and wildtype littermates show no difference in size or expression level of Celsr3 or Celsr1 protein. Brain lysate from *Celsr3*^*-/-*^ mutants is included as a control for antibody specificity. **(B, C)** DiI labeling of open book preparations shows that commissural axons from *Celsr3*^*R1548Q*^ mutants (n=6) **(C)** exhibit AP randomization after crossing the floor plate, similar to *DG*^*F/-*^*;Sox2*^*Cre*^, *ISPD*^*L79*/L79\**^, and *Celsr3*^*-/-*^ mice. Scale bar = 50μm. **(D)** Quantification of open book preparations, with the number of injection sites showing anterior turning over the total number of injection sites indicated above each bar. *p< 0.001, Student’s T-test. **(E-J)** L1 immunohistochemistry **(E, F)** and DiI labeling of thalamocortical **(F, I)** and corticothalamic **(G, J)** axons show no defects in internal capsule formation in *Celsr3*^*R1548Q*^ mutants. Scale bar = 500μm. **(K)** Proposed model for Dystroglycan:Celsr3 interactions in guiding commissural axons.

We first examined spinal commissural axon crossing and anterior turning in *Celsr3*^*R1548Q/R1548Q*^ mutants in open-book preparations. Remarkably, post-crossing commissural axons exhibited randomization along the AP axis, similar to *Celsr3*^*-/-*^, *ISPD*^*L79*/L79\**^, and *DG*^*F/-*^*;Sox2*^*Cre*^ mutants (Figure 6B). Quantification shows that only 10/48 injection sites in *Celsr3*^*R1548Q/R1548Q*^ mutants had normal anterior turning, while the remaining 38 exhibited AP randomization. Importantly, *Celsr3*^*R1548Q/R1548Q*^ mutants did not exhibit the floorplate stalling phenotypes that are seen in *ISPD*^*L79\**^, *DG*^*F/-*^*;Sox2*^*Cre*^, and *Slit/Robo* compound mutants. These results suggest that Celsr3 interacts with Dystroglycan through its LG1 domain to direct the proper anterior turning of post-crossing commissural axons.

In contrast to the results we observed in the spinal cord, the internal capsule and other axon tracts in the forebrains of *Celsr3*^*R1548Q/R1548Q*^ mutants appeared normal by both immunostaining and DiI labeling (Figure 6E-J). Therefore, the requirement for Dystroglycan:Celsr3 interactions appears to be context dependent, and the defects in internal capsule formation in *Dystroglycan* and *ISPD*^*L79*/L79\**^ mutants likely reflect Dystroglycan interactions with other LG-domain containing proteins such as Laminins or Slits.

## Discussion

Severe forms of Dystroglycanopathy (WWS, MEB) are characterized by profound neurodevelopmental defects that can include type II lissencephaly, hydrocephalus, hindbrain hypoplasia, and defects in white matter. Interestingly, congenital mirror movements, which arise from improper decussation of descending corticospinal axons as they pass through the brainstem, have been reported in isolated cases of dystroglycanopathy, suggesting that axon tract abnormalities may contribute to this disorder (Ardicli et al., 2017; Longman et al., 2003). Using a model of severe dystroglycanopathy (*ISPD*^*L79\**^) and *Dystroglycan* conditional mutants, we now show that Dystroglycan is required for formation of several major axon tracts in the forebrain. Using conditional deletion of *Dystroglycan* and *DG*^*βcyto/-*^ mice, we find that Dystroglycan functions non-cell autonomously as an extracellular scaffold to guide axons within the brain and spinal cord. Furthermore, we identify the transmembrane axon guidance receptor Celsr3 as a novel binding partner for Dystroglycan, and show that this interaction is mediated by a conserved region in the LG1 domain of Celsr3. We show that the Dystroglycan:Celsr3 interaction is context dependent *in vivo*, and is required for the correct anterior turning of post-crossing commissural axons. Taken with our previous results, these findings demonstrate that axon guidance defects are a key feature of dystroglycanopathy, which arise due to Dystroglycan’s interaction with multiple ECM proteins, secreted axon guidance cues, and transmembrane axon guidance receptors.

### Dystroglycan regulates several aspects of nervous system development by binding to multiple proteins

The neurological abnormalities in patients with dystroglycanopathy are extremely heterogeneous, ranging from mild cognitive defects to severe and widespread structural abnormalities (Godfrey et al., 2011). Mouse models of dystroglycanopathy and conditional *Dystroglycan* knockouts have demonstrated that Dystroglycan is required for multiple aspects of neuronal development, including neuronal migration, axon guidance, synapse formation, glial development, and maintenance of the blood-brain barrier (Clements et al., 2017; Fruh et al., 2016; McClenahan et al., 2016; Menezes et al., 2014; Michele et al., 2002; Moore et al., 2002; Myshrall et al., 2012; Saito et al., 2003; Satz et al., 2008; Satz et al., 2010; Wright et al., 2012). The widespread nature of these defects reflects the reiterative function of Dystroglycan throughout neurodevelopment and its interactions with multiple partners.

During early neurodevelopment, Dystroglycan binding to basement membrane proteins (Laminin, Perlecan) maintains the attachment of neuroepithelial cells that serve as scaffolds for neuronal migration in the brain and retina. The subsequent role for Dystroglycan in regulating axon guidance reflects its interactions with multiple binding partners. Dystroglycan organizes ECM proteins in the basement membrane as a permissive growth substrate, restricts the extracellular localization of the secreted cue Slit, and interacts with the transmembrane receptor Celsr3. Deletion of *Dystroglycan* selectively from neurons or deletion of its intracellular signaling domain does not affect either neuronal migration or axon guidance, suggesting that it functions exclusively non-cell autonomously in the neuroepithelium as an extracellular scaffold in these contexts (Clements et al., 2017; Satz et al., 2010).

At later stages of neurodevelopment, Dystroglycan is expressed in neurons, where it regulates specific subsets of perisomatic inhibitory synapses and hippocampal LTP (Fruh et al., 2016; Satz et al., 2010; Zaccaria et al., 2001). It is unclear which proteins Dystroglycan interacts with at synapses, although Neurexins are possible candidates, as they have been shown to bind to Dystroglycan through their LG domains (Reissner et al., 2014; Sugita et al., 2001a). Several other LG-domain containing proteins that contain the conserved Dystroglycan binding motif (Celsrs, CNTNAPs, Thrombospondins, Laminins) are also localized to synapses, suggesting that Dystroglycan may have a complex and context specific role in synapse formation and maintenance. Importantly, patients with milder forms of dystroglycanopathy can have cognitive defects even in the absence of any obvious structural abnormalities in the brain, which may reflect the role of synaptic Dystroglycan.

### Dystroglycan:Celsr3 interactions during axon guidance

In this study, we identify a novel interaction between Dystroglycan and the transmembrane receptor Celsr3. Remarkably, when we disrupted this interaction *in vivo* by mutating a key residue in a highly conserved region of the LG1 domain in Celsr3 (*Celsr3*^*R1548Q*^), we observed AP randomization of post-crossing spinal commissural axons *in vivo,* similar to *ISPD*^*L79*/L79\**^, *DG*^*F/-*^*;Sox2*^*Cre*^, and *Celsr3*^*-/-*^ mutants. Based on our data that Dystroglycan functions non-cell autonomously (Figure 1F, G) and does not require its intracellular domain (Figure 1E, G), and previous work showing that Celsr3 functions cell autonomously in commissural axons (Onishi et al., 2013), we propose a model in which Celsr3 in the growth cones of commissural axons must bind in *trans* to Dystroglycan in the neuroepithelium and/or basement membrane as axons cross the floorplate (Figure 6F). Previous work has shown that Celsr3 and Fzd3 are required to direct the anterior turning of commissural axons in response to a Wnt gradient in the floor plate. How this occurs is still not well understood, but it involves downstream signaling pathways involving Jnk and atypical PKC (Lyuksyutova et al., 2003; Onishi et al., 2013; Onishi and Zou, 2017; Wolf et al., 2008).

Despite the similarities in internal capsule phenotypes in *ISPD*^*L79*/L79\**^, *DG*^*F/-*^*;Sox2*^*Cre*^, and *Celsr3*^*-/-*^ mutants, disrupting the Dystroglycan:Celsr3 interaction did not affect formation of this axon tract in *Celsr3*^*R1548Q*^ mutants. During internal capsule formation, Celsr3 is required in Isl1^+^ neurons in the prethalamus and ventral telencephalon to form a pioneering axon scaffold that is required for TCAs and CTAs to cross the DTB (Feng et al., 2016). Our data suggest that this occurs independent of Celsr3 interactions with Dystroglycan. It is possible that the cadherin repeats of Celsr3 mediate homophilic interactions between the Isl1^+^ axons, but this has not been tested directly. Celsr3 and Fzd3 are also required cell autonomously for peripheral extension of motor axons into the limb, where they form a complex with Ephrin-A2, -A5, and Ret (Chai et al., 2014). In contrast, *Dystroglycan* and *Celsr3*^*R1548Q*^ mutants do not display any defects in peripheral motor axon growth (data not shown). Therefore, the importance of Dystroglycan:Celsr3 interactions during axon guidance are context dependent.

If Dystroglycan:Celsr3 interactions are not required for internal capsule formation, what could explain the severe forebrain axon guidance phenotypes in *ISPD*^*L79*/L79\**^ and *DG*^*F/-*^*;Sox2*^*Cre*^ mutants? Dystroglycan has the ability to bind multiple LG-domain containing partners simultaneously through its extensive glycan chains, making it difficult to ascribe function to a single molecular interaction. However, Slits are likely candidates in the forebrain, as *Slit* and *Robo* mutants exhibit defects in internal capsule, anterior commissure and lateral olfactory tract development similar to *ISPD*^*L79*/L79\**^ and *DG*^*F/-*^*;Sox2*^*Cre*^ mutants (Bagri et al., 2002; Bielle et al., 2011; Fouquet et al., 2007; Lopez-Bendito et al., 2007). In addition to directly repelling axons, Slits also regulates the migration of cells that function as intermediate guideposts for the axons that form the internal capsule and lateral olfactory tract, suggesting the axon guidance phenotypes may be secondary to neuronal migration defects (Bielle et al., 2011; Fouquet et al., 2007). Dystroglycan may influence neuronal migration and axon guidance in the forebrain by regulating the distribution of Slit proteins, similar to its role in the ventral midline of the spinal cord. Determining precisely how these pathways interact to regulate axon tract formation will require careful spatial and temporal manipulation of their expression *in vivo*.

In light of Dystroglycan’s binding to multiple ligands important for nervous system development, it is interesting to note that Dystroglycan displays differential glycosylation patterns in muscle, glia, and even between neuronal subtypes. For example, glial Dystroglycan migrates at ∼120kD, Dystroglycan in cortical/hippocampal neurons migrates slightly higher (∼140kD), whereas Dystroglycan in cerebellar Purkinje neurons migrates at ∼180kD (Satz et al., 2010). How these differences in glycosylation arise and whether they affect binding to different ligands in distinct cell types remains unclear.

### Evolutionary conservation of Dystroglycan function during axon guidance

Dystroglycan and its binding partners have evolutionarily conserved roles in regulating axon guidance. The *C. elegans* Dystroglycan homologue DGN-1 is required for follower axons to faithfully track along pioneer axons (Johnson and Kramer, 2012). Similarly, FMI-1, the *C. elegans* homologue of Celsr3, is involved in both pioneer and follower axon guidance in the ventral nerve cord (Steimel et al., 2010). FMI-1 phenotypes could be rescued by expressing the regions encompassing either the cadherin repeats or the EGF and LG domains of FMI-1, suggesting that it may function in both a homophilic and heterophilic manner, depending on the context. In *D. melanogaster*, the Dystroglycan homologue Dg functions in both neurons and glial cells to guide the proper targeting of photoreceptor axons to the optic lobe (Shcherbata et al., 2007). *Slit* mutants and RNAi to *Robo1/2/3* show a remarkably similar photoreceptor targeting phenotype to *Dg* mutants, which arises from a failure to form a proper boundary between the lamina glia and the lobula cortex (Tayler et al., 2004). *Flamingo*, the Drosophila homologue of Celsr3, functions at a subsequent step in visual system development to non-cell autonomously regulate synaptic choice of photoreceptors (Chen and Clandinin, 2008; Lee et al., 2003; Senti et al., 2003). The remarkable similarities in axon targeting defects seen in *Dystroglycan*, *Slit* and *Celsr3* mutants across species suggests that their interactions are evolutionarily conserved.

In summary, our results establish a widespread role for Dystroglycan in regulating axon tract formation during neurodevelopment. We also identify Celsr3 as a novel binding partner for Dystroglycan and find that their interaction is required for anterior turning of post-crossing commissural axons. By functioning as an extracellular scaffold that binds multiple ECM proteins, secreted axon guidance cues, and transmembrane receptors, Dystroglycan plays a critical role in many aspects of neural circuit development and function.

## Acknowledgments

We thank members of the Wright laboratory for their assistance and discussion throughout the course of this study; Marc Freeman, Kelly Monk, Tianyi Mao, Martin Riccomagno and Randal Hand for comments on the manuscript; Krissy Lyons, Jessica Barowski, and Kylee Rosette for technical assistance. We thank David Ginty, in whose lab this work was started, for advice and financial support during the initial phases of the work and for generation of *Celsr3*^*R1548Q*^ mice, and Kevin Campbell for providing *DG*^*βcyto*^ mice. This work was supported by NIH grant NS091027 (KMW), the Medical Research Foundation of Oregon (KMW), ARC convention number 17/22–079 (FT), and startup funds from Vollum Institute/OHSU (KW).

## Author Contributions

Conceptualization: K.M.W.; Methodology: C.G. and K.M.W.; Investigation: B.L., N.P., and K.M.W.; Writing-original draft: K.M.W.; Funding Acquisition: K.M.W.; Supervision: F.T. and K.M.W.

## Star Methods

### Generation and analysis of mutant mice

*ISPD*^*L79*\*^ (Wright et al., 2012), *Dystroglycan*^*Flox/Flox*^ (Michele et al., 2002), *Dystroglycan*^*βcyto*^ (Satz et al., 2009), *Sox2*^*Cre*^ (Hayashi et al., 2002), *FoxG1*^*Cre*^ (Hebert and McConnell, 2000), *Gbx2*^*CreERT2*^ (Chen et al., 2009) *Emx1*^*Cre*^ (Gorski et al., 2002), *Dlx5/6*^*Cre*^ (Stenman et al., 2003), and *Wnt1*^*Cre*^ mice were maintained on a C57Bl/6J background. *Ai9/R26*^*LSL-TdTomato*^ mice were maintained on an outbred CD1 background.

*Celsr3*^*R1548Q*^ mice were generated using CRISPR/Cas9 pronuclear injection by the HHMI/Janelia Farm Gene Targeting and Transgenic Facility. The gRNA (5′-AAAAGTCATGCTTCTCGTTC-3′) was co-injected with 163 bp ssDNA (5’-ATGTCTGATCCTAATGGTCCCACTCCACTTCACTCAGGTTTGCAACTGTGCA ACCCAGCGGGCTACTCTTCTACAACGGG<u>CAG</u>CTGAACGAGAAGCATGACTT TTTGGCTCTAGAGCTTGTGGCTGGCCAAGTGCGGCTTACATATTCCACGGGTGGGTGCTC-3’) and Cas9 protein with a concentration of 5:25:25 ng/ul. Eight founders from 58 pups were identified with mutations in Celsr3. Correctly targeted mutations were then confirmed by PCR, followed by Sanger sequencing. Three correctly targeted *Celsr3*^*R1548Q*/+^ founders were obtained, and after outcrossing to the outbred CD1 strain for two generations, *Celsr3*^*R1548Q/+*^ mice were intercrossed to generate *Celsr3*^*R1548Q/R1548Q*^ homozygous mutants. *Celsr3*^*R1548Q*^ mice were genotyped using the following primers: Fwd: 5’-CACTGGCATCTCCCACACTA-3’ and Rev: 5’-GGGACACCTGAGAGGATTCA-3’. PCR products were then incubated with PvuII, which cuts the CAGCTG site generated in the *Celsr3*^*R1548Q*^ mutants.

Mice were handled and bred in accordance with the Oregon Health and Science University IACUC guidelines. Embryos were obtained from timed pregnancies, with the date of plug appearance counted as e0.5. To generate *Dystroglycan* conditional knockouts, *Dystroglycan*^*+/-*^; *Cre*^*+*^ male breeders were crossed to *Dystroglycan*^*Flox/Flox*^ females. All conditional knockout analyses used *Dystroglycan*^*F/+*^; *Cre*^*+*^ littermates. Phenotypic analysis was conducted on at least five different offspring obtained from at least three different litters, using at least two different male breeders, without regard to sex of animals analyzed. Mice were genotyped by PCR as previously described.

### Immunohistochemistry and anterograde tract tracing

For analysis of brains, P0 mice were euthanized by decapitation, brains were removed and fixed in 4% paraformaldehyde at 4° overnight. For L1 immunostaining, brains were washed three times for 30 minutes each in PBS, then embedded in low melt agarose. 150 μm thick vibrotome sections were collected and washed once in PBS, blocked for 30 minutes in PBS + 0.25% TritonX-100, 5% goat serum, then incubated in primary antibody diluted in blocking buffer at 4° for two days. Sections were washed in PBS five times for thirty minutes each, then incubated in secondary antibody diluted in blocking buffer at room temperature, overnight. Sections were then washed five times for 1 hour each in PBS, with DAPI (1:5000) included in the second wash step. Sections were then mounted on Permafrost slides, light protected with Fluoromount-G (Southern Biotech), and imaged. For anterograde tract tracing, DiI crystals were inserted into the cortex or thalamus of brains, returned to 4% paraformaldehyde, and incubated at 4° for 5-7 days. Brains were then embedded in low melt agarose, 150 μm thick vibrotome sections were collected in PBS, incubated in DAPI (1:5000) for 30 minutes, washed once in PBS for five minutes, mounted, and imaged on a Zeiss M2 Imager equipped with ApoTome. Images were processed in Zeiss Zen Blue and Adobe Photoshop 6 software.

### Open book preparations

Embryos were collected at E12.5 and fixed for 30 minutes in 0.5% paraformaldehyde. Spinal cords were then removed, split along the roof plate, the meninges were removed, and the flattened spinal cords were fixed in 4% paraformaldehyde for four hours at room temperature. DiI crystals were then inserted along the lateral margin of the spinal cord and tissue was incubated in 4% paraformaldehyde at room temperature overnight. Open book preparations were then imaged on a Zeiss ZoomV-16 dissecting microscope at 50X magnification.

### Binding assays

293T cells were transfected with constructs encoding Fc-tagged Dystroglycan (DG-Fc), AP-tag alone, AP-Celsr3-LG1, AP-Celsr3-LG-2, AP-Celsr3-LG1^R1548Q^, or AP-Slit-Cterm. After recovery, cells were maintained in OptiMEM for 48-72 hours, after which supernatant was collected, concentrated by centrifugation (Amicon, 10kD molecular weight cutoff), and exchanged to binding buffer (20mM Hepes, pH 7.0, 150mM NaCl, 2.5 mM CaCl_2_). DG-Fc was coupled to Protein-A agarose beads for 6 hours at 4°, beads were washed once in binding buffer, and 5nM of AP-tagged ligand was added and beads were incubated at 4° overnight, rocking.

For endogenous Dystroglycan binding assays, brains from postnatal day 7 (P7) mice were homogenized in a 10X volume of PBS + 1% Triton, incubated for 1 hour at 4°, rocking and insoluble material was removed by centrifugation at 3400xg. Supernatant was incubated with WGA-agarose beads at 4° overnight, then competed off the WGA-beads with 500mM N-acetyl-D-glucosamine, followed by dialysis in binding buffer at 4° overnight. WGA-enriched lysate was then incubated with AP-tagged ligands (5nM) pre-coupled to NiNTA beads.

For all binding experiments, beads were washed five times with binding buffer to remove unbound material. Bound proteins were eluted by boiling in 1X LDS sample buffer with 50mM DTT for 10 minutes, resolved by SDS-PAGE, transferred to PVDF membranes, blocked for 60 minutes in 5% nonfat milk in TBS + 0.1% Tween-20 (TBST), then probed with antibodies diluted in blocking buffer at 4° overnight. Membranes were washed three times for 10 minutes in TBST, incubated with secondary antibody diluted in blocking buffer with 5% nonfat milk, washed three times for 10 minutes in TBST, and developed with SuperSignal ECL Pico.

For live cell binding assays, Cos7 cells plated on poly-D-lysine were transfected with myc-tagged full-length Dystroglycan. 48 hours after transfection, cells were incubated with AP-tagged ligand at 37° for 30 minutes. Cells were then washed five times in HBSS, fixed with 4% paraformaldehyde + 60% acetone for 30 seconds, and washed five times in HBSS. Plates were then incubated at 67° for 90 minutes to inactivate endogenous peroxidase activity. Cells were washed twice in AP buffer (100mM Tris, pH 9.0, 50mM MgCl2), then incubated with BCIP/NBT in AP buffer for 30-60 minutes until signal developed. The AP reaction was stopped by washing cells twice in HBSS + 50mM EDTA, and cells were imaged on a Zeiss ZoomV-16 dissecting microscope at 100X magnification.

### Celsr3^R1548Q^-GFP generation and *<u>in vitro</u>* <u>assays</u>

Celsr3^R1548Q^-GFP was generated by QuickChange Mutagenesis from the parent Celsr3-GFP vector (Chai et al., 2014). 293T cells grown on PDL-coated coverslips were transfected with either Celsr3-GFP or Celsr3^R1548Q^-GFP, and analyzed 48 hours later. For analysis of protein localization by immunocytochemistry, cells were briefly fixed in 4% PKS (paraformaldehyde in Krebs + sucrose) for 30 minutes at room temperature. Cells were then washed three times in PBS for 10 minutes, blocked for 30 minutes in PBS + 0.25% TritonX-100, 5% goat serum, then incubated in primary antibody diluted in blocking buffer at 4° overnight. Cells were washed in PBS five times for five minutes each, then incubated in secondary antibody diluted in blocking buffer at room temperature for two hours. Cells were then washed five times for 5 minutes each in PBS, with DAPI (1:5000) included in the second wash step. Coverslips were then mounted and imaged.

For total and cell surface expression, 293T cells in 60mm plates were transfected with either Celsr3-GFP or Celsr3^R1548Q^-GFP and allowed to recover for 48 hours. Cell surface labeling was done with the Pierce Cell Surface Protein Isolation Kit, according to the manufacturer’s instructions.

### Quantification and statistical analysis

No statistical methods were used to predetermine sample sizes, but they were similar to our previous work (Clements et al., 2017; Wright et al., 2012). For all phenotypic analyses, tissue was collected from at least five different offspring obtained from at least three different litters, using at least two different male breeders. For open book preparations, each injection site was scored blindly as to whether it exhibited normal anterior turning or AP randomization by three lab members. All analysis was done blind to genotype. Data was tested for normality and statistical analysis was conducting using JMP Pro version 13.0 (SAS Institute). Comparison between two groups was analyzed using a Student’s t test; Comparison between two or more groups was analyzed using a one-way ANOVA and Tukey’s post hoc test. *p<0.0001.

---

**Supplemental Figure1:**
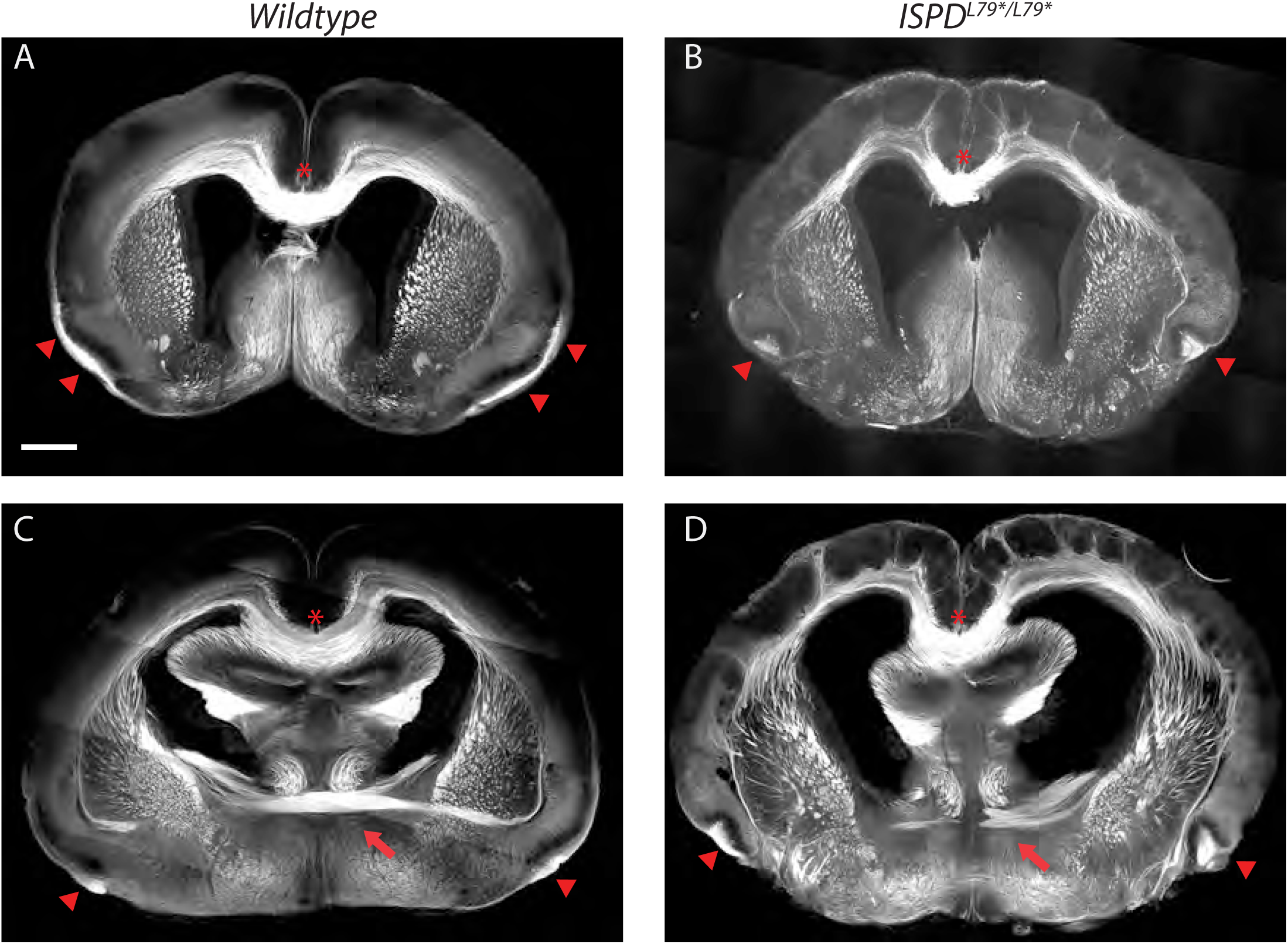
Anterior commissure, lateral olfactory tract and corpus callosum phenotypes in *ISPD*^*L79*/L79\**^ mutants. **(A-D)** L1 staining was used to label forebrain axon tracts in wildtype **(A, C)** and *ISPD*^*L79*/L79\**^ mutants **(B, D)**. The corpus callosum (red asterisk) in *ISPD*^*L79*/L79\**^ mutants appears largely normal compared to controls. In contrast, the anterior commissure (red arrows) is thinner and disorganized in *ISPD*^*L79*/L79\**^ mutants (D). The lateral olfactory tract (red arrowheads) extends along the pial surface of the ventrolateral telencephalon in controls **(A, C)**, whereas it appears hyperfasciculated and projects deeper into the piriform cortex as a disorganized bundle in *ISPD*^*L79*/L79\**^ mutants **(B, D)**. Scale bar = 500μm

**Supplemental Figure2:**
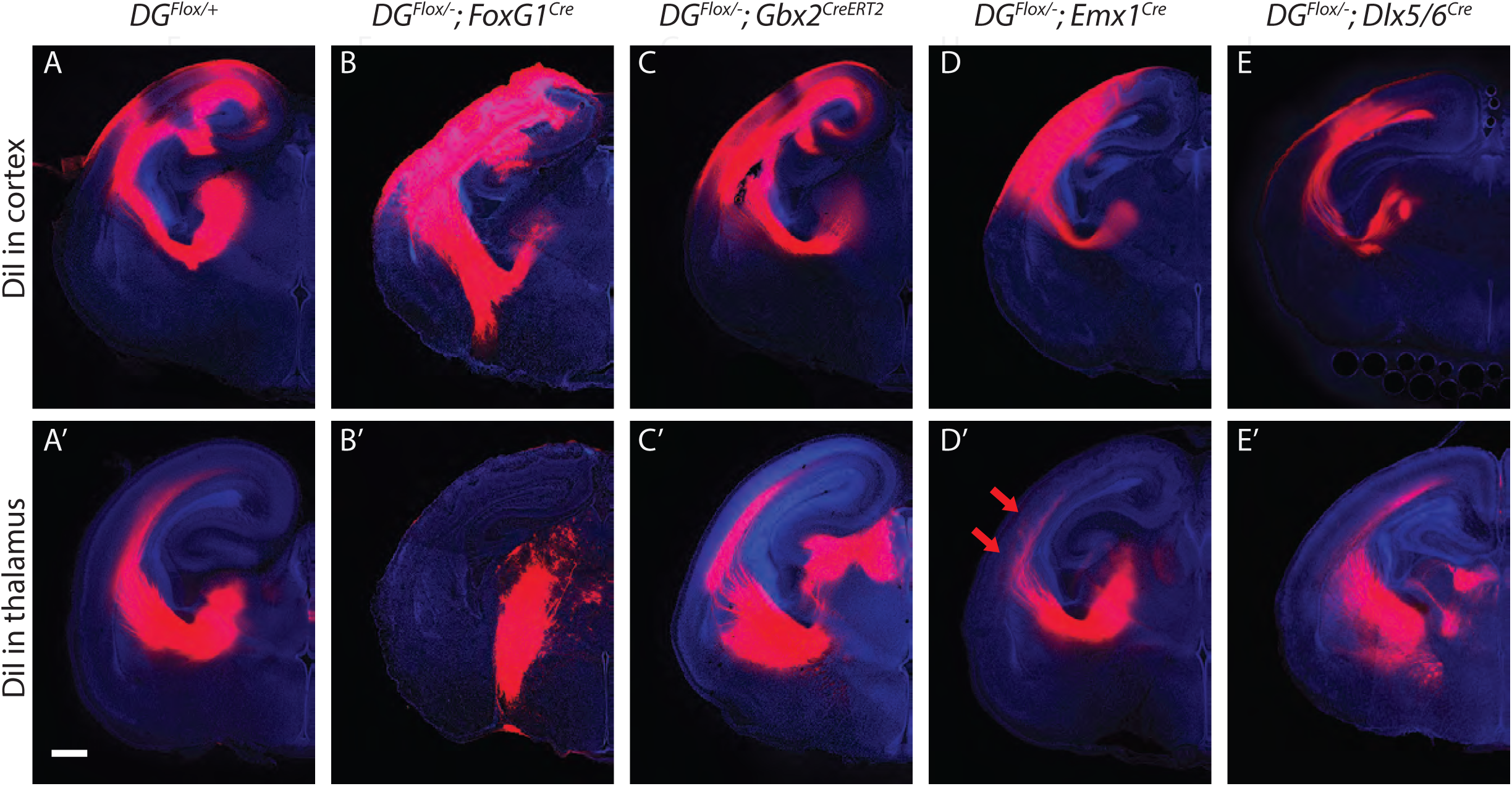
DiI labeling of CTAs and TCAs in *Dystroglycan* conditional mutants. **(A-E)** DiI injections into the cortex (top row) or thalamus (bottom row) of *Dystroglycan* conditional mutants labels CTAs and TCAs, respectively. CTAs **(B)** in *DG*^*F/-*^*;FoxG1*^*Cr*e^ mutants take an abnormal trajectory through the ventral telencephalon, and TCAs **(B’)** fail to cross the DTB and instead extend ventrally out of the diencephalon. CTAs in *DG*^*F/-*^*;Gbx2*^*Cr*eERT2^ **(C)**, *DG*^*F/-*^*;Emx1*^*Cr*e^ **(D)** in *DG*^*F/-*^*;Dlx5/6*^*Cr*e^ **(E)** mutants are normal, as are TCAs in *DG*^*F/-*^*;Gbx2*^*Cr*eERT2^ **(C’)**, *and DG*^*F/-*^*;Dlx5/6*^*Cr*e^ **(E’)** mutants. TCAs in *DG*^*F/-*^*;Emx1*^*Cr*e^ **(D’)** mutants project through the internal capsule normally, but project into the upper layers prematurely upon entering the cortex (red arrows). Scale bar = 500μm.

**Supplemental Figure3:**
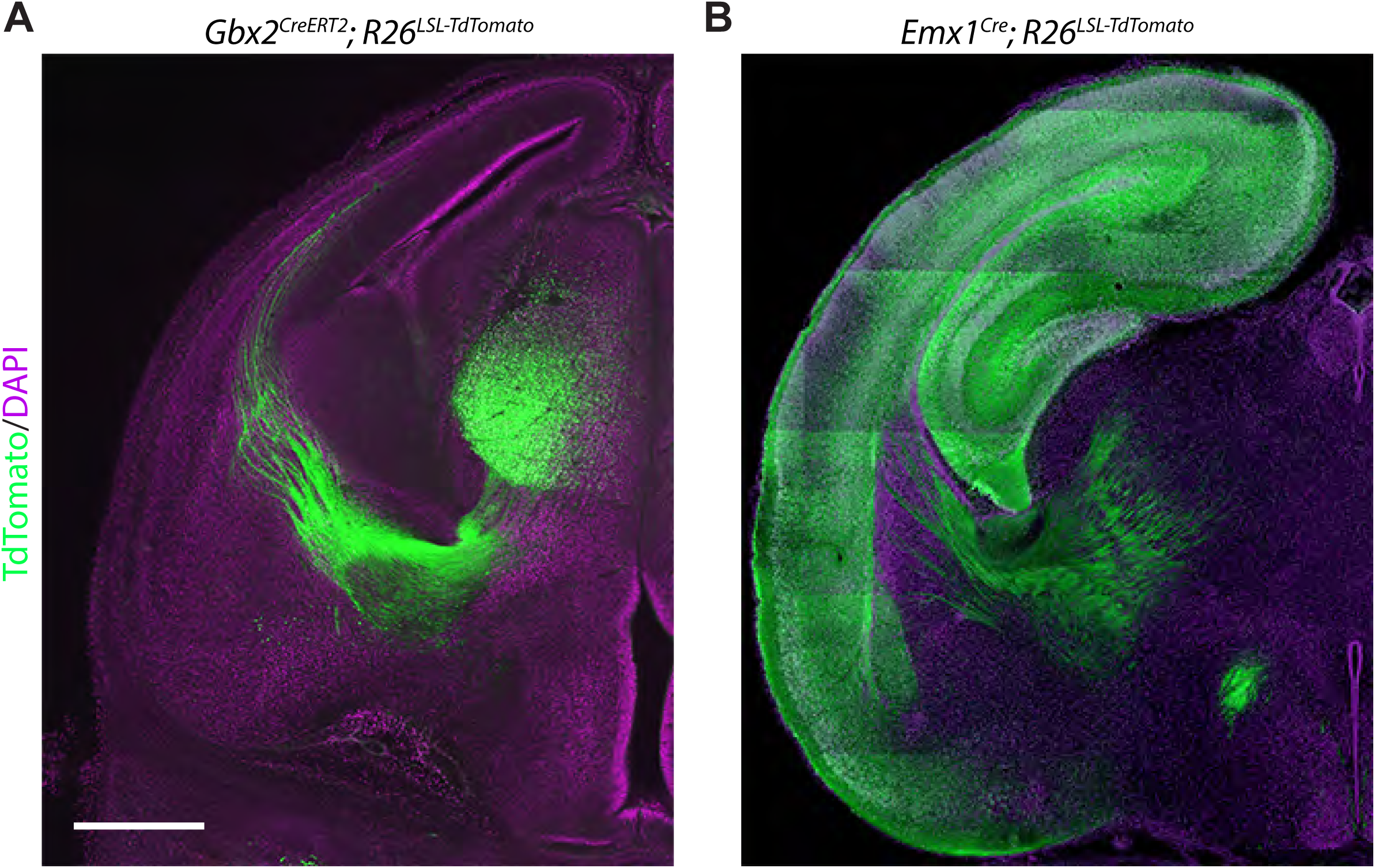
Recombination pattern in *Gbx2*^*CreERT2*^ and *Emx1*^*Cre*^ mice. **(A)** *Gbx2*^*CreERT2*^ mice crossed to the *AI9; Rosa26*^*lox-stop-lox-tdTomato*^ reporter were dosed with 2.5mg tamoxifen at e10.5. Analysis of brains at E16 showed recombination of the *tdTomato* reporter (green) in thalamic neurons/axons. **(B)** *Emx1*^*Cre*^ mice crossed to the *AI9; Rosa26*^*lox-stop-lox-tdTomato*^ reporter showed recombination of the *tdTomato* reporter (green) in cortical neurons/axons at P0. Scale bar = 500μm.

**Supplemental Figure4:**
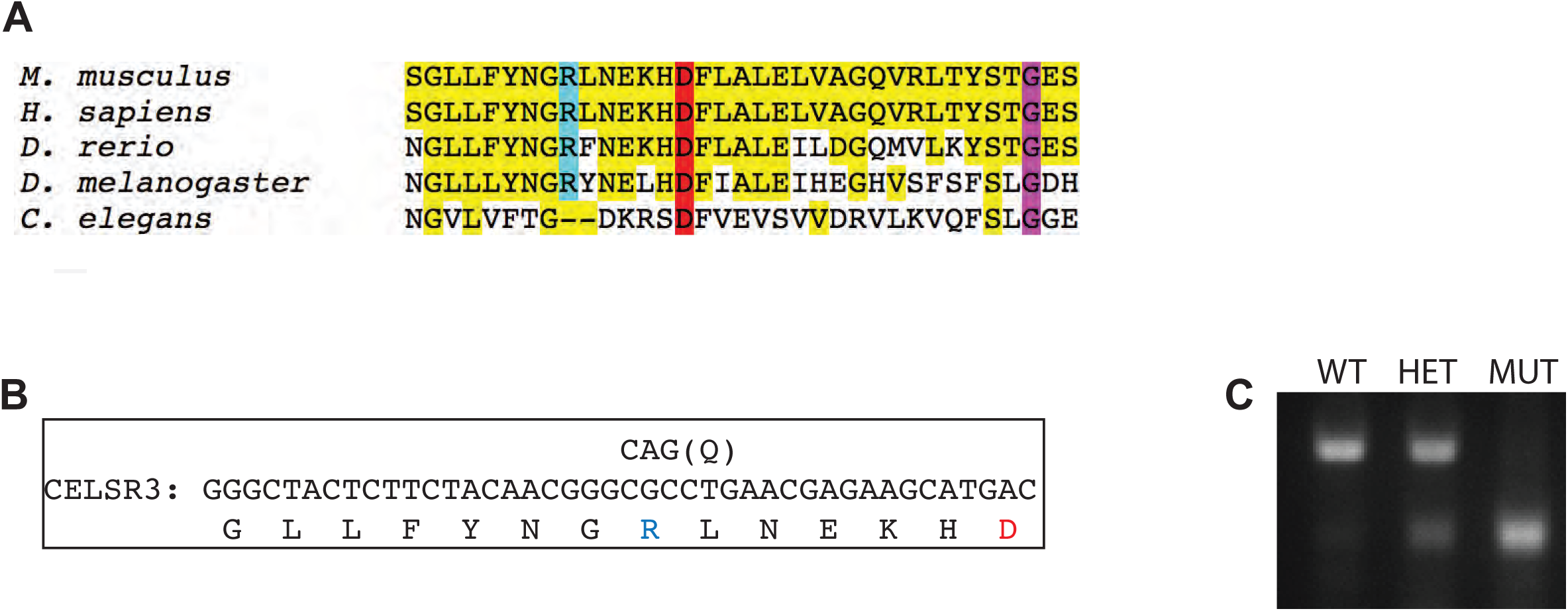
Analysis of *Celsr*^*R1548Q*^ mutants. **(A)** Evolutionary conservation of Celsr3-LG1 region that forms a putative binding interface with Dystroglycan. **(B)** Schematic of the Celsr3 nucleotide and amino acid sequence highlighting the specific sequence targeted to generate *Celsr*^*R1548Q*^ knock-in mice. **(C)** Genotyping of wildtype, *Celsr*^*R1548Q/+*^, and *Celsr*^*R1548Q/R1548Q*^ mutants.

